# The possible anti-inflammatory activity of macrolide antibiotics in male albino rat models

**DOI:** 10.1101/2020.10.20.340950

**Authors:** Mohamed Hamed, Esmail Abdelmonem, Mahmoud Zayed, Sameh Shaban, Iman El Khashab, Soheir Abu-El-Azm

**Author notes:** Corresponding author, Soheir Abu-El-Azm; Mohamed Hamed.

## Abstract

Recently, there has been increasing evidence on the use of macrolide antibiotics in treatment of chronic inflammatory diseases through mechanisms distinct from their antibacterial activity. The key desired effect lies somewhere between these two therapeutic potentials and has not been identified yet. The aim of the present study was to evaluate the anti-inflammatory activity of the macrolide antibiotic azithromycin in formaldehyde induced arthritis and carrageenan induced air pouch in albino rats in comparison to the anti-rheumatic drug meloxicam. Results of Formaldehyde induced arthritis revealed that pretreatment of animals with a single daily dose of either azithromycin (10, 20 and 40 mg/kg) or meloxicam (4 mg/kg) for 15 days produced a significant reduction in inflammation size. Histopathological study showed that Formaldehyde produced marked inflammatory cell infiltration, congestion of blood vessels and soft tissue edema which were attenuated by azithromycin in dose dependent manner. The radiological study revealed that azithromycin attenuates soft tissue edema, periarticular bone resorption, narrowing of joint spaces and joint deformities induced by formaldehyde. This effect was marked with 40mg/kg azithromycin pre-treatment. In carrageenan, induced air pouch, results demonstrate that group of animals pretreated with azithromycin (10, 20 and 40 mg/kg) or meloxicam (4 mg/kg) for 6 days significantly attenuated the mean increase in total leukocyte count in air pouch exudate. In conclusion, the present work showed that azithromycin has antiinflammatory activity in the models tested and suggests that it can exert therapeutics effects independent of its anti-bacterial activity. Also, the anti-inflammatory effect of azithromycin was potent and even comparable to that of meloxicam.

## Introduction

Inflammation is one of the most basic processes that occur in human’s body. It is a complex innate defensive response of the body to tissue injury, trauma, infection or exposure to radiation, thereby decreasing damage of tissues (Hunag et al., 2004; Cuzzocrea, 2005). It can be divided into 2 types: acute and chronic inflammations (Anon, 2011). Causes of inflammation are infections initiated by bacteria, viruses, bacteria toxins, fungus, yeast, autoimmune diseases such as arthritis, chemicals as acids and bases and exposure to radiation such as ultraviolet radiation (Chen et al., 2018). Inflammation involves a series of events that are initiated by inflammatory mediators such as histamine, eicosanides, complement, cytokines, serotonins, bradykinins, prostaglandins, Tumor Necrosis Factor (TNF), Platelet Activating Factor (PAF) and interleukins (Yao and Narumiya, 2019). This is followed by the inflammatory response that occurs in the form of blood vessel dilatation, increase vascular permeability, release of inflammatory mediators at the injury site, phagocytosis by neutrophils, macrophages and other immune cells (Cuzzocrea, 2005). New anti-inflammatory drugs are discovered and developed based on their effects on signal transduction as anti-cytokine agents to control diseases where cytokines and other non-prostaglandin components of neurodegenerative diseases and chronic inflammatory manifest (Rao et al., 2010; Patil et al., 2019; Zappavigna et al., 2020).

Macrolide antibiotics are defined for their broad-spectrum activity that makes them a drug of choice for treatment of conditions caused by a wide spectrum of bacterial infections. They are derived from Streptomyces species. A macrolytic lactone ring is very common in macrolides in which sugars (one or more) can be attached (Vázquez-Laslop and Mankin, 2018). Members of macrolide antibiotics can be differentiated from each other by their chemical substitutions through their structure in different atoms of amino, carbon and neutral sugars (Bayarski, 2006). The macrolides antimicrobial mechanism of action is based on inhibition of protein synthesis by reversibly binding bacterial ribosome 50s subunit thus inhibiting peptidyl-tRNA translocation. The inflammation mechanism is adopted by many methods as inhibition of T-cell proliferation (Lee et al., 2011; Kannan et al., 2014). One of the main characteristics that cause them to achieve their effects, is their ability to accumulate within leukocytes and then transported into the site of infection (Čulićet al., 2001). Macrolides can be used to treat respiratory tract infections (sinusitis, pharyngitis, lower respiratory tract infections), skin and soft tissue infections, mycobacterial infections, sexually transmitted diseases, cervicitis, urethritis, H. pylori infections (clarithromycin, as part of triple therapy), inflammations (bronchitis, arthritis) (Romero et al., 2017; Kelly et al., 2018). The anti-inflammatory activity of some macrolides like tacrolimus (FK506) is proved to be effective as an immunosuppressive drug mainly in cases of organ transplantation (Campagne et al., 2019). Macrolides can be used as alternative treatment for penicillins and cephalosporins in allergic patients. Several studies have assured the presence of an anti-inflammatory action exerted by macrolide antibiotics. It was proved that macrolides can inhibit peripheral blood mononuclear cells proliferation in vitro thus leading to decrease in the amount of superoxide which is formed by neutrophils (Konno et al., 1993).

The objective of this study is to test and assess the anti-inflammatory activity of macrolide antibiotics. The aim of the work is to evaluate the difference and strength of the antiinflammatory effect of different concentrations of the macrolide antibiotic azithromycin in inflammation models. Our findings suggest a promising role of macrolide antibiotic azithromycin in management of inflammation independent of its antimicrobial activity.

## Materials and methods

### Drugs and chemicals

Azithromycin in powder form was obtained as parenteral infusion product (Zithromax 500 mg; Pfizer, USA). Azithromycin was prepared by dissolving the powder in 5 ml of distilled water and then diluting the resultant solution by 20 ml of distilled water. Meloxicam in solution form was obtained as intramuscular ampoules of the product (Melocam 15 mg/ml; Amooun pharmaceuticals, Egypt). Carrageenan (pure λ+κ carrageenan powder) was obtained from Sigma Aldrich (Germany) and was freshly prepared before administration by dissolving in 100% saline.

### Animals

Male Albino rats weighing 180-200 g were used. The rats were housed in cages (6 in each) at ordinary room temperature, exposed to natural daily light/dark cycle, fed on standard laboratory chow, tab water and labtum. The rats were acclimatized for one week and randomly allocated into groups.

### Formaldehyde induced arthritis in rats (Selve, 1949)

Chronic soft tissue inflammation and arthritis were induced in the male Albino rats by injection of 0.1 ml of 2% formaldehyde solution into the plantar region of the left hind paw of the rats on day 1 and day 3 of the experiment which lasted for a period of fifteen days.

Forty-two rats were divided into seven groups; each consist of six rats. Group I (Normal control): rats were injected intra peritoneal by 0.2 ml distilled water. Group II (Formaldehyde induced): rats were injected intraperitoneal by 0.2 ml distilled water one hour before the injection of sub plantar 0.1 ml of formaldehyde 2%. Group III: rats were given intraperitoneal Azithromycin 10 mg/kg one hour before the injection of sub plantar 0.1 ml of formaldehyde 2%. Group IV: rats were given intraperitoneal Azithromycin 20mg/kg one hour before the injection of sub plantar 0.1 ml of formaldehyde 2%. Group V: rats were given intraperitoneal Azithromycin 40mg/kg one hour before the injection of sub plantar 0.1 ml of formaldehyde 2%. Group VI: rats were given intraperitoneal Meloxicam 4mg/kg one hour before the injection of sub plantar 0.1 ml of formaldehyde 2%. Group VII: rats were given intraperitoneal Azithromycin 10 mg/kg plus Meloxicam 4 mg/kg one hour before the injection of sub plantar 0.1 ml of formaldehyde 2%. Drugs were given intraperitoneal in a single daily dose for whole length of the experiment.

### Carrageenan induced air pouch model (Edwards et al., 1981)

Air pouch was produced by subcutaneous injection of 20 ml of air into the intra-scapular area of the back on day one and the air pouch was then maintained by re-inflation with 10 mL of air three days later. On day six, 0.1 ml carrageenan 1% suspension was injected directly into air pouches of lightly restrained (hand held), conscious animals. The animals were sacrificed by cervical decapitation 4 hours after carrageenan injection and exudate was lavage out of the pouch under direct visualization. Forty-two were divided into seven groups, each contains six rats. Group I (normal control): rats were injected by 0.1 mL saline intra pouch and were given 2 mL distilled water intraperitoneal. Group II (carrageenan induced): rats were injected by 0.1 mL of 1% carrageenan suspension intra pouch and were given 2 mL distilled water intraperitoneal. Group III: rats were injected by 0.1 mL of 1% carrageenan suspension intra pouch and were given Azithromycin 10 mg/kg intraperitoneal. Group IV: rats were injected by 0.1 mL of 1% carrageenan suspension intra pouch and were given Azithromycin 20 mg/kg intraperitoneal. Group V: rats were injected by 0.1 mL of 1% carrageenan suspension intra pouch and were given Azithromycin 40mg/kg intraperitoneal. Group VI: rats were injected by 0.1 mL of 1% carrageenan suspension intra pouch and were given Meloxicam 4mg/kg intraperitoneal. Group VII: rats were injected by 0.1 mL of 1% carrageenan suspension intra pouch and were given Azithromycin 10mg/kg plus Meloxicam 4 mg/kg intraperitoneal. Drugs were given for 6 days after air pouch induction intraperitoneal in a single daily dose. The last dose was given 1 hour before carrageenan injection.

### X-ray and histopathological examination

At the end of the experimental design on the 15^th^ day, the left hind paw was imaged antero-posteriorly by X-ray. The rats of all groups were sacrificed and their left hind paw was excised just above the ankle joint for weight measurement and histopathological examination. The weights of left hind paws of all groups were measured for comparison. Autopsy samples were taken from the paw in different groups and fixed in 10% formol saline for twenty-four hours. Washing was done in tap water then serial dilutions of alcohol (methyl, ethyl and absolute ethyl) were used for dehydration. Specimens were cleared in xylene and embedded in paraffin at 56 degree in hot air oven for twenty-four hours. Paraffin bees wax tissue blocks were prepared for sectioning at 4 microns by slidge microtome. The obtained tissue sections were collected on glass slides, de-paraffinized and stained by hematoxylin and eosin stain, then examined through Zeiss Axiolab A1 light microscope (Oberkochen, Germany; Banchroft et al; 1996).

### Total Leukocyte Count (TLC)

At the end of experiment on day six, the exudate was collected and examined for Total Leukocyte Count (TLC) where the cells were counted in a standard hemocytometer chamber.

### Statistical analysis

The significance of differences between control and treated groups in terms of mean increase in weight of left hind paw (Formaldehyde-induced model), Total Leukocyte Count (carrageenan induced air pouch model) and percent inhibition were evaluated by oneway analysis of variance (ANOVA) followed by the Turkey-Kramer comparison test. All statistical analyses were done using the program GraphPad Prism 5. The level of significance was set at P<0.05 in all cases.

## Results

### Meloxicam reduced formaldehyde induced arthritis in rats

Injection of formaldehyde (0.1 ml of 1% solution) into the planter surface of the hind paw of the rats on day 1 and day 3 produced marked increase in paw weight until day 15 (p<0.05) the mean weight was 2.68 ± 0.14 (g) vs. 1.44 ± 0.04 for control (Table 1, Fig. 1a and c-f). Administration of Azithromycin intraperitoneal in single doses of 10, 20 and 40 mg/kg for the whole length of the experiment 1 hour before formaldehyde treatment, significantly attenuated the increase in paw weight with percentages inhibition of 32.46, 34.30 and 40.11 % respectively (Table 1, Fig 1b). Animal groups pretreated with meloxicam 4 mg/kg significantly attenuated the increase in paw weight with percentages inhibition of 29.30 % (Table 1, Fig 1b). In addition, combination of treatment (Azithromycin 10 mg/kg and Meloxicam 4 mg/kg) produced 32.40 % inhibition.

**Figure 1:**
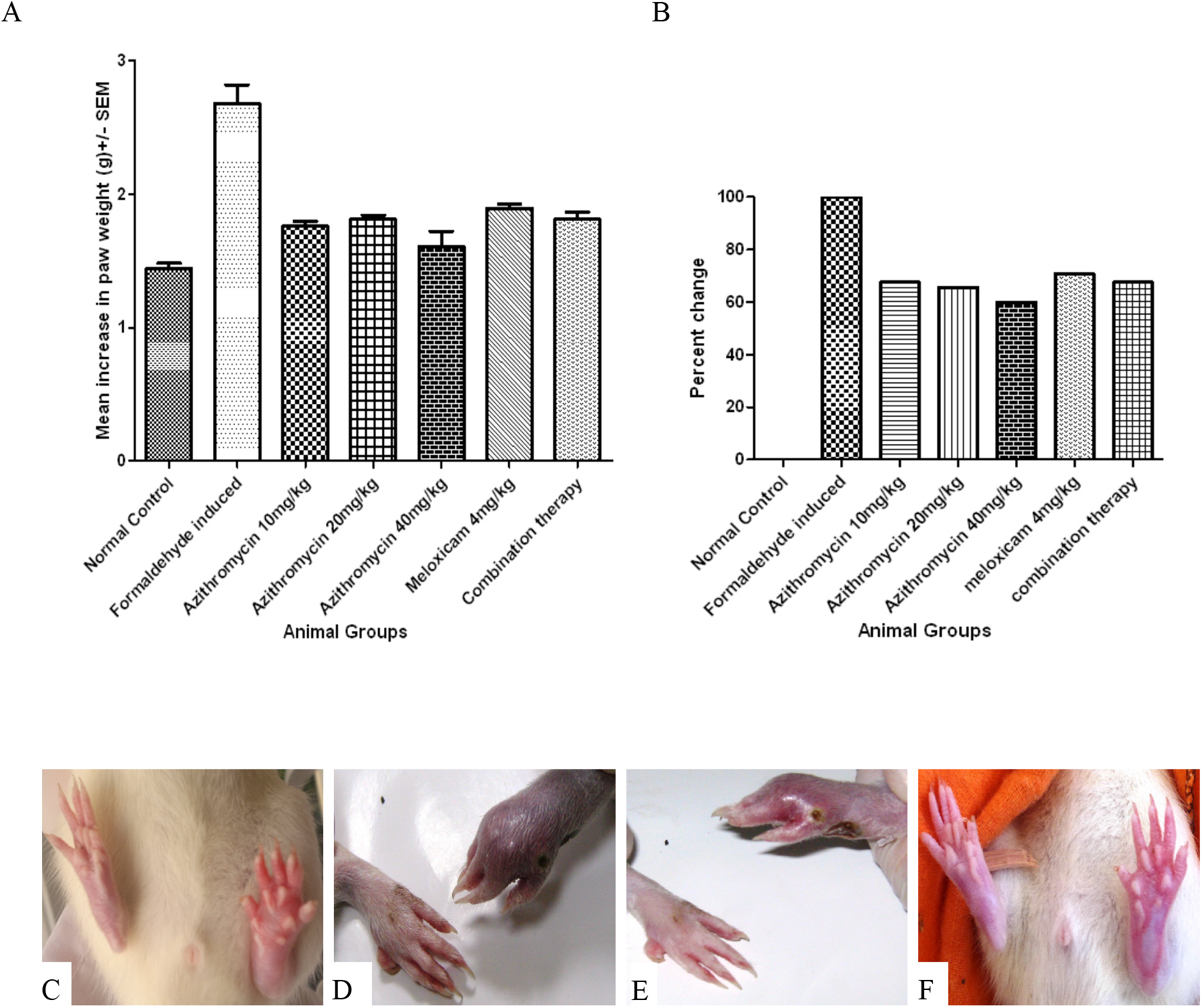
Azithromycin reduces inflammation size in a comparable manner to meloxicam. **A)** Change in inflammation size determined as means increase in paw weight in groups treated with azithromycin (10, 20, 40mg/kg), Meloxicam (4mg/kg) and their combination following formaldehyde induced arthritis. Standard error mean (SEM) of three independent measurements in indicated. **B)** Percent change of paw weight shown in (**A**) normalized to the normal control group. **C)** Rat hind paw show effect of formaldehyde (group 2) vs. normal control. **D)** Rat hind paw of azithromycin (40mg/kg) vs. formaldehyde group. **E)** Rat hind paw of meloxicam (4mg/kg) vs. formaldehyde group. **F)** Rat hind paw treated with combination therapy vs. normal (group 1).

**Table 1:**
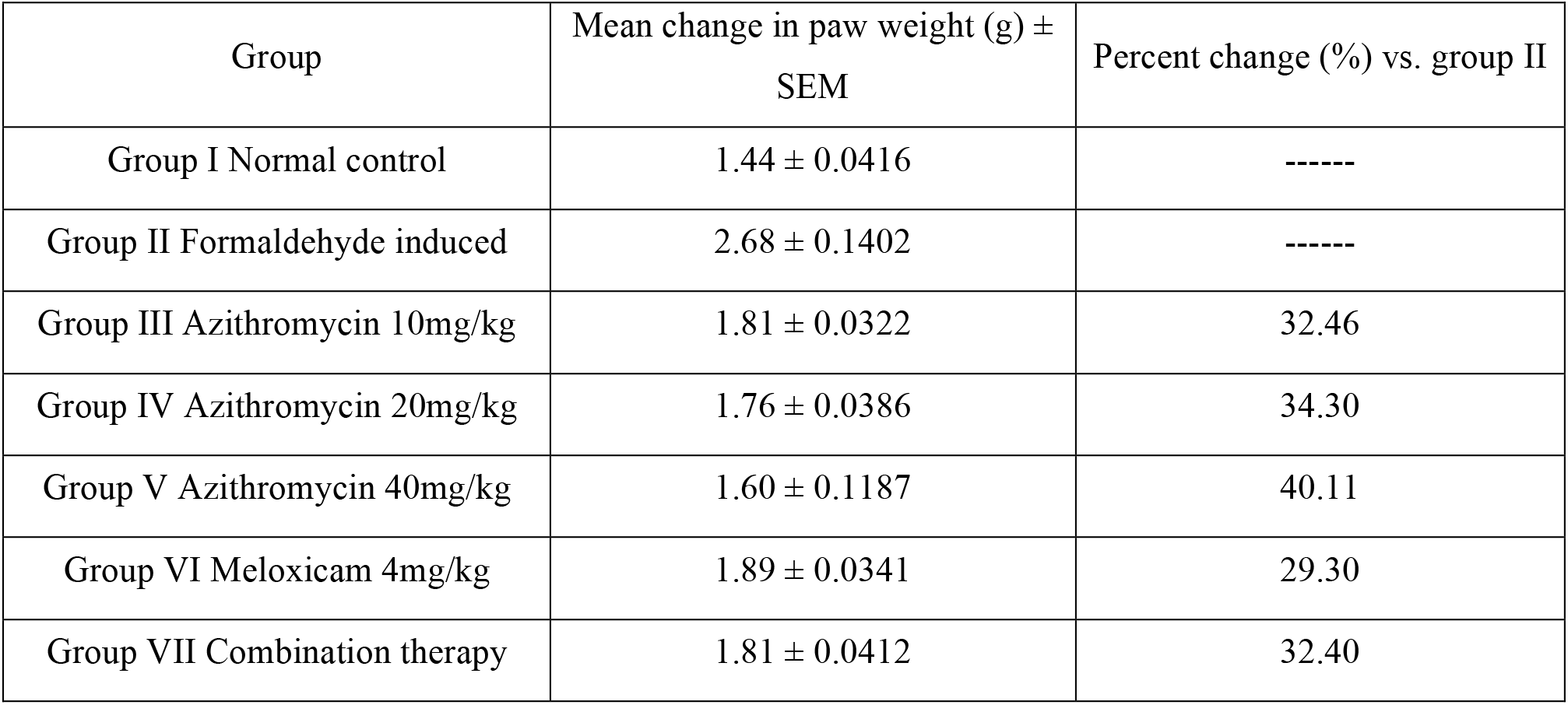
Effect of Azithromycin (10, 20 and 40 mg/kg), Meloxicam (4 mg/kg) and their combination on formaldehyde induced arthritis in Male Albino rats

| Group | Mean change in paw weight (g) ± SEM | Percent change (%) vs. group II |
| --- | --- | --- |
| Group I Normal control | 1.44 ± 0.0416 | ----- |
| Group II Formaldehyde induced | 2.68 ± 0.1402 | ----- |
| Group III Azithromycin 10mg/kg | 1.81 ± 0.0322 | 32.46 |
| Group IV Azithromycin 20mg/kg | 1.76 ± 0.0386 | 34.30 |
| Group V Azithromycin 40mg/kg | 1.60 ± 0.1187 | 40.11 |
| Group VI Meloxicam 4mg/kg | 1.89 ± 0.0341 | 29.30 |
| Group VII Combination therapy | 1.81 ± 0.0412 | 32.40 |

Injection of formaldehyde produced marked paw tissue inflammation that involved all layers in form of acanthosis, necrosis and degeneration in the prickle cells of the epidermis associated with inflammatory cell infiltration, dilation in blood vessels in both dermal and subcutaneous tissue (Fig 2b). This histopathological study also showed thickening with inflammatory cell infiltration in periosteum surrounding contact resorbed bone (Fig 2c) vs. control (Fig 2a). Administration of azithromycin 10, 20 and 40 mg/kg/day intraperitoneal for the whole length of the experiment attenuated these signs of inflammation (Fig 2d, e and f). In addition, the inflammatory cell infiltration was markedly attenuated in azithromycin treated (40mg/kg) and meloxicam (4mg/kg) and their combination (Fig 2g, h and i).

**Figure 2:**
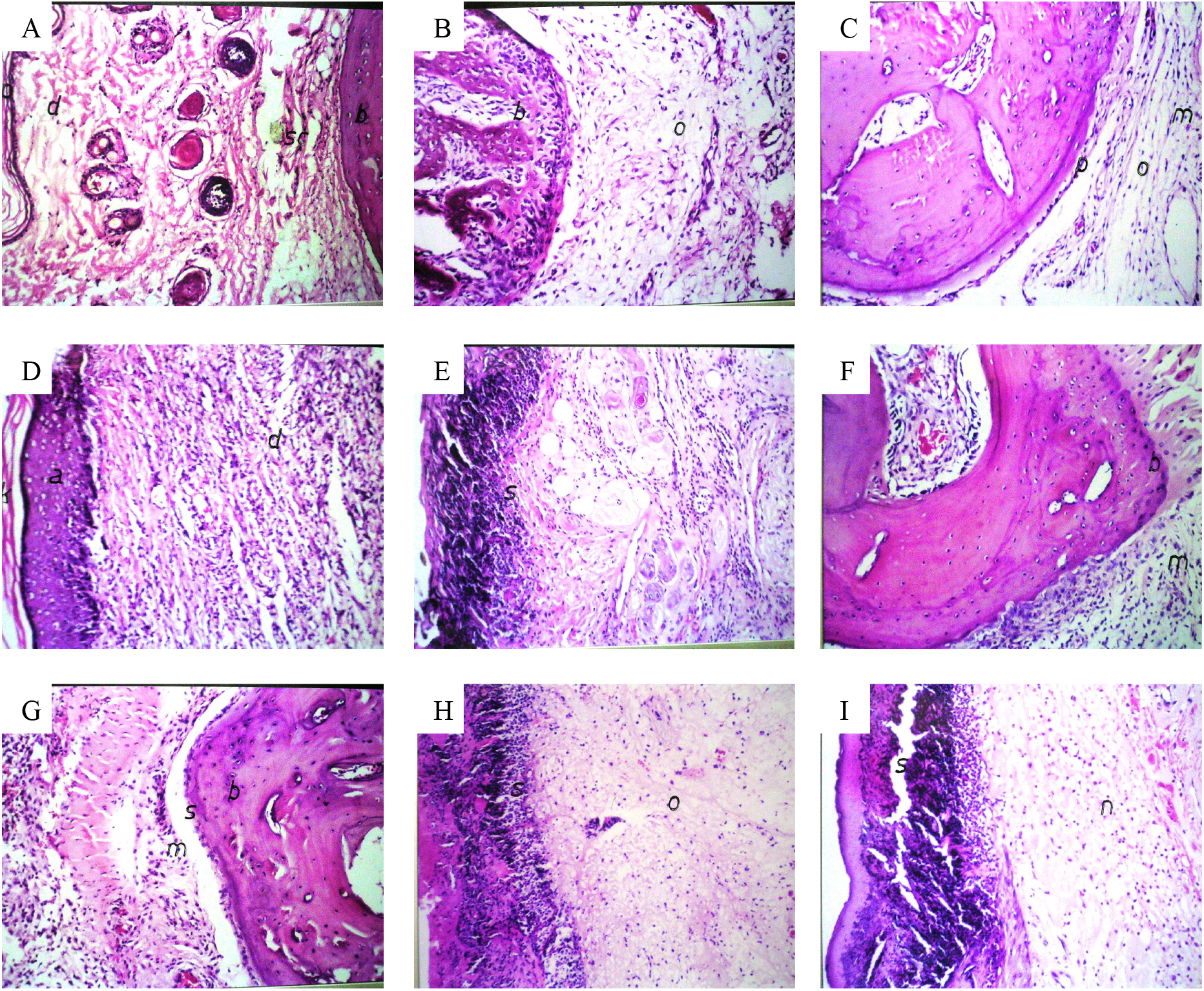
Azithromycin reduces edema inflammatory cell infiltration of rat hind paw. **A)** paw of rat in group 1 showing normal histological structure of the epidermis (p), dermis (d), subcutaneous tissue (Sc) and bone (b), H *f* E x 40. **B)** paw of rat in group 2 showing acanthosis of the prickle cell layer in the dermis (a) with inflammatory cell infiltration in the dermis (d), H *f* E x 40. **C)** paw of rat in group 2 showing thickening with inflammatory cells infiltration in periosteum (s) surrounding the contact resorbed bone (b), H *f* E x 64. **D)** paw of rat in azithromycin 10mg/kg (group 3) showing moderate bone resorbtion (b) with moderate inflammatory cell infiltration (o), H *f* E x 40. **E)** paw of rat in azithromycin 20mg/kg (group 4) showing hyalinization of the dermal tissue (h) underneath epidermis (s) with moderate inflammatory cell infiltration, H *f* E x 40. **F)** paw of rat in azithromycin 40mg/kg (group 5) showing few inflammatory cell infiltration (o) in dermis underneath epidermis (s), H *f* E x 40. **G)** paw of rat in azithromycin 40mg/kg (group 5) showing thick periosteum (p), less edema (o) and few inflammatory cell infiltration (m) in surrounding subcutaneous tissue, H *f* E x 40. **H)** paw of rat in meloxicam 4mg/kg (group 6) showing inflammatory cell infiltration (m) surrounding normal bone structure, H *f* E x 40. **I)** paw of rat in combination group (azithromycin 10mg/kg and meloxicam 4mg/kg) (group 7) showing epidermis (s) with few inflammatory cell infiltration (n), H *f* E x 64.

Analysis of joint radiographs at the termination of the experiment revealed that injection of formaldehyde produced evident radiological manifestations of inflammation and arthritis in the form of marked soft tissue edema, periarticular bone resorption, narrowing of the joint spaces and joint deformities (Fig 3c and d) vs. control (Fig 3a and b). Groups treated with azithromycin 10, 20 and 40 mg/kg/day, meloxicam 4 mg/kg/day and their combination showed significant attenuation of these signs of inflammation by decreasing soft tissue edema, joint deformity and destruction (Fig 3c-g). This effect was significant with azithromycin 40 mg/kg and meloxicam 4 mg/kg (Fig 3e, f and g).

**Figure 3:**
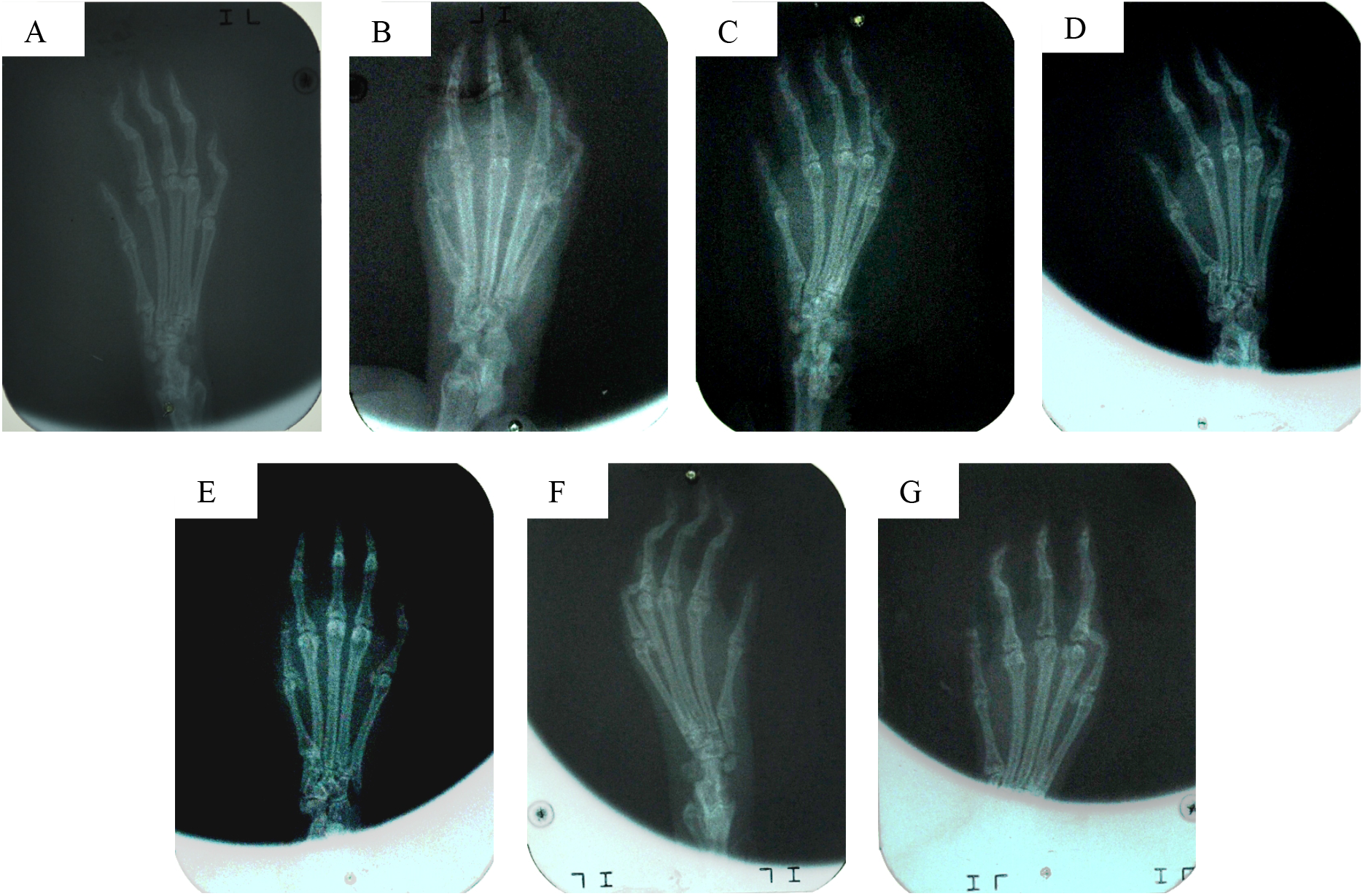
Azithromycin reduces joint deformities. **A)** Photo (IV a) Rat of normal control group (group I). **B)** Photo (IV b) Rat of formaldehyde induced arthritis (group 2). **C)** Photo (V a) Rat treated with azithromycin (10 mg/kg/day) (group 3). **D)** Photo (V b) Rat treated with azithromycin (20 mg/kg/day) (group 4). **E)** Photo (VI a) Rat treated with azithromycin (40 mg/kg/day) (group 5). **F)** Photo (VI b) Rat treated with meloxicam (4 mg/kg/day) (group 6). **G)** Photo (VI c) Rat treated with a combination therapy of azithromycin (10 mg/kg/day) and meloxicam (4 mg/kg/day) (group 7).

### Total leucocytes count is affected by meloxicam treatment in carrageenan induced air pouch model

Carrageenan injection (0.1 ml of 1% suspension) into air pouch (group 2) produced a significant increase in air pouch leucocytes count. The mean TLC in group 2 was 2140 ± 66.33 / cmm Vs 880 ± 0.80/ cmm in controlled group (group 1) (Table 2, Fig 4a). Group of animals pretreated with azithromycin (10, 20 and 40 mg/kg/day) or meloxicam (4 mg/kg/day) for 6 days significantly attenuated the mean increase in TLC with percentages inhibition by 59.3%, 66.3%, 77.4 and 67.7 % respectively (Table 2, Fig 4b). Groups of animal treated with azithromycin (10 mg/kg/day) and meloxicam (4 mg/kg/day) showed significant reduction of TLC by 68.8 % (Fig 4b).

**Figure 4:**
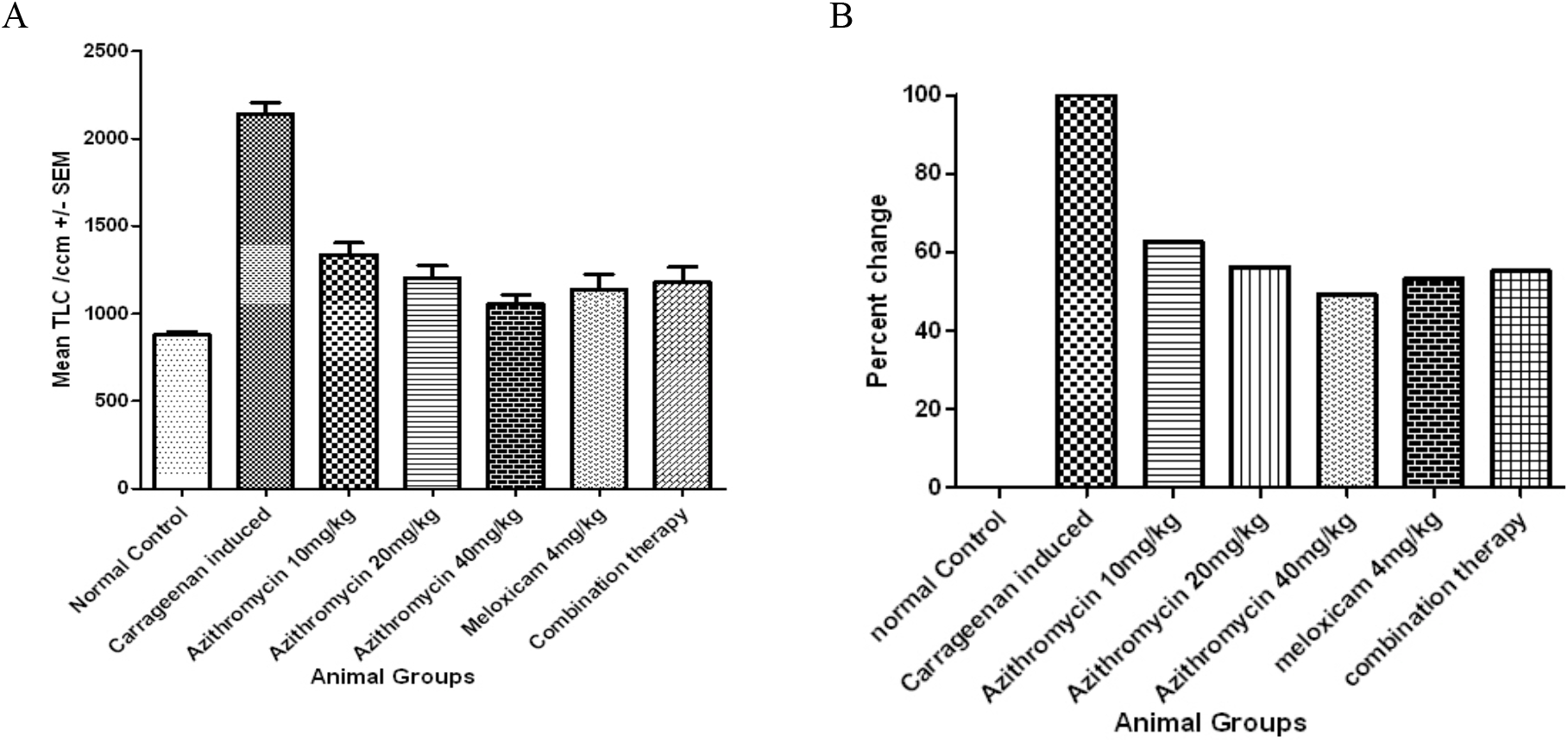
Azithromycin TLC in carrageenan induced air pouch model. **A)** Effect of Azithromycin (10, 20, 40mg/kg), Meloxicam (4mg/kg) and their combination on TLC in carrageenan induced air pouch model. Standard error mean (SEM) of three independent measurements in indicated. **B)** Percent change of paw weight shown in (**A**) normalized to the normal control group.

**Table 2:** Effect of Azithromycin (10, 20 and 40 mg/kg), Meloxicam (4 mg/kg) and their combination on TLC in carrageenan induced air pouch model

| Group | Mean TLC/cmm ± SEM | Percent change (%) vs. group II |
| --- | --- | --- |
| Group I Normal control | 880 ± 8.80 | ----- |
| Group II Positive induced | 2140 ± 66.33 | ----- |
| Group III Azithromycin 10mg/kg | 1337 ± 69.55 | 59.33 |
| Group IV Azithromycin 20mg/kg | 1203 ± 71.45 | 66.33 |
| Group V Azithromycin 40mg/kg | 1053 ± 54.81 | 77.44 |
| Group VI Meloxicam 4mg/kg | 1140 ± 83.91 | 67.78 |
| Group VII Combination therapy | 1180 ± 86.56 | 68.89 |

These results indicate a role of azithromycin in the reduction of inflammation at doses lower than those required for antimicrobial activity and comparable to anti-inflammatory agents such as meloxicam.

## Discussion

Inflammation is a complicated local response to any stimuli or certain types of auto-immune diseases as arthritis and bronchitis. It involves a series of reactions that involve several inflammatory mediators as cytokines, neutrophils, adhesion molecules, complement, and IgG (Trowbridge and Emling, 1997). Abnormal inflammatory cell accumulation is a character of the majority of inflammatory diseases whether they originate from infective or non-infective disease (Yao and Narumiya, 2019). Macrolide is a class of antibiotics that have been used since long time ago. It plays an important role in the treatment of different types of infectious diseases. One of the characteristic of this family is possibility that the intracellular accumulation of the drug which alters the functions of the host cell. This character has recently elevated a new interest in the possible therapeutic effects of macrolides in clinical settings other than infections. Based on these findings, the present work was designed to evaluate the antiinflammatory activity of azithromycin in some experimental models of inflammation (Vázquez-Laslop and Mankin, 2018).

In formaldehyde-induced arthritis model, results revealed that Injection of formaldehyde (0.1 ml of 1% solution) into the planter surface of the hind paw of the rats on day 1 and day 3 produced marked increase in paw weight until days 15 (p<0.05) the mean increase was 2.68 ± 0.14 (g) vs 1.44 ± 0.04 for control. Administration of Azithromycin I.P in a single dose 10, 20, 40 mg/kg/day for the whole length of the experiment starting 1 hour before Formaldehyde significantly attenuated the increase in paw weight with percentage of inhibition of 32.46, 34.30 and 40.11 % respectively. Animal group pretreated with meloxicam 4 mg/kg significantly attenuated the increase in paw weight with percentage of inhibition of 29.30 %. In addition, combination of treatment (Azithromycin 10 mg/kg + Meloxicam 4 mg/kg) produced 32.40 % inhibition. Scaglione and Rossoni (1998) studied the anti-inflammatory effects of roxithromycin, azithromycin and clarithromycin against nimesulide. They concluded that all three macrolides tested were able to reduce the inflammation caused by carrageenan in the rat paw model only slightly less than nimesulide. Ianaro and collagues (2000) studied the effect of roxithromycin, clarithromycin, erythromycin and azithromycin on the generation of some mediators and cytokines involved in the inflammatory process induced by carrageenan in the rat pleurisy model. It was found that exudate volume and leukocyte accumulation were both dose-dependently reduced by these drugs. Furthermore, they significantly reduced PGE2, nitrate and TNF-α level in pleural exudate. Ren et al. (2004) demonstrated that Erythromycin suppressed weak debris-induced osteoclastic bone resorption by inhibiting inflammatory osteoclastogenesis through down-regulation of NF-KB signaling pathway. Therefore, macrolide antibiotic represents a promising therapeutic candidate for the prevention and treatment of aseptic loosening in patients with total joint arthroplasts. Li et al. (2006) suggested that EM703, a new 12-membered macrolide derivative of erythromycin without antibacterial effects, reduced bleomycin induced pulmonary fibrosis in mice by anti-inflammatory action through regulation of transforming growth factor beta (TGF-B) signaling in lung fibroblasts.

In the present study, the degree of inflammation induced by Carrageenan, as measured by total leukocyte count in air pouch exudates, was determined in different animal groups. Pretreatment with Azithromycin (10, 20 and 40 mg/kg/day) on meloxicam (4 mg/kg/day) for 6 days or their combination, significantly attenuated the increase in TLC produced by Carrageenan with percentages inhibition of 59.33, 66.33, 77.44, 67.78 and 68.89 respectively. It was observed that the effect of 40 mg/kg was compatible to meloxicam and there is no additional antiinflammatory in the combination groups. Khobragade et al. (2010), reported that Roxithromycin 20 mg/kg and Erythromycin 40 mg/kg have anti-inflammatory activity in Carrageenan-induced rat paw edema model also Macrolide exhibited less anti-inflammatory activity than ibuprofen (100 mg/kg), and there is no additional anti-inflammatory activity in the combination group. There are some studies which stated that macrolide antibiotics possess antiinflammatory action. It was proved that macrolides can inhibit peripheral blood mononuclear cells proliferation in vitro. That can lead to decrease the amount of superoxide which is formed by neutrophils and also modify cytokines release (Konno et al., 1993). Some Animal models were used in vivo in order to estimate macrolide anti-inflammatory action in the airways. Inside a rat tracheal mucosa roxithromycin and erythromycin both can inhibit microvascular leakage as well as neutrophil recruitment in action to intravenous lipopolysaccharide (Tamaoki et al., 2000). It was proved that FK506 which is a macrolide with anti-inflammatory (but not antibacterial action) can be used in order to manage rheumatoid arthritis. FK506 has the ability to increase patients’ activity through muscle strength improvement and decreasing the inflammation as well as pain. It suppresses NFAT localization which causes T-cells inactivation through working as an immunosuppressive drug which then causes inhibition in calcineurin. It also acts through T-cells cytokines inhibition and the TNF-α and IL-1β (derived inflammatory cytokines antigen presenting cell) (Sasakawa et al., 2004; Majithia and Geraci, 2007). Bronchitis which is simply inflammation that affects mucous membranes of the bronchi which cause problem in the airways leading to airflow limitation from the trachea into the lungs (Sevilla-Sánchez et al., 2010). According to Čulić et al. (2001), some macrolides as Roxithromycin has been reported to exert some antiasthmatic effects, based on inhibition of bronchial hyperresponsiveness and polymorphonuclear leukocyte superoxide production. The anti-inflammatory bronchial has also been reported in a clinical trial.

To address the question of whether there is anti-inflammatory effect of macrolide or not. The effect of macrolide were studied in patients which do not take oral corticosteroids. However still there is macrolides possibility like erythromycin which can give their effect through inhibiting inhaled corticosteroid metabolism by their action on one of the P450 cytochrome enzymes (CYP 3A4). On the other hand, there are some new macrolides like azithromycin and roxithromycin which have no effect or little one on the CYP 3A4. 12 weeks of roxithromycin treatment in a study of patient with asthma showed decrease in there bronchial hyperresponsiveness. Several reports suggested that the use of roxithromycin might be considered very useful asthma treatment, however in the double-blind absence and in placebo controlled studies that cannot be confounded through macrolide effect on steroid metabolism, assuming that macrolide can be used in asthma treatment is still premature (Shimizu et al., 2004). Kelly et al. (2018) described that use of macrolide therapy, especially azithromycin, over a long-time course resulted in reduction of exacerbations frequency in patients.

The subcellular mechanisms of the anti-inflammatory effects of macrolides are still unknown. The fact that there are several distinct inflammatory pathways and that each of which proceeds via a cascade of biological events, makes it difficult to identify which of these events is the target of macrolides and the definite mechanism of their anti-inflammatory action. Čulić et al. (2002) hypothesized that acute neutrophilic degranulation and oxidative burst may contribute to an antimicrobial effect and delayed responses may contribute to an anti-inflammatory effect of azithromycin. It was reported that macrolide antibiotics inhibit NO generation from fibroblasts. They speculated that this may be one of the mechanisms leading to the favorable modification of airway inflammation with macrolide therapy (Terao et al., 2003). Ou et al (2007) speculated that the inhibitory effect of roxithromycin on airway mucus hyersecretion may be mediated through reduction of NF-KB activation, neutrophil infiltration and release of inflammatory cytokines in the lung. Gao et al (2010) reported that inhibition of cytokine production may result from a modification of gene expression and reduction in mRNA expression and protein release for cytokines (Kannan et al., 2014). Apart from antibacterial activity, macrolides are documented to have anti-inflammatory actions, and the same has been documented in animal models, (Ianaro et al., 2000) as well as in clinical scenarios, (Wallwork et al., 2006). The anti-inflammatory activity of macrolides is proposed to be independent of (cox-2) inhibition (Vázquez-Laslop and Mankin, 2108). In addition, a study by Porter et al. (2016) showed that macrolide anti-inflammatory activity might contribute to macrolide antiviral activity. The anti-inflammatory properties of macrolides form the basis for their experimental use in bronchial asthma and air way inflammatory and infective conditions (Maselli et al., 2011; Vaz et al., 2011). Since Meloxicam is known inhibitor of cox-2, the combination of two anti-inflammatory drug acting via different mechanisms should have caused an increased effect also as seen in Khobragade et al. (2011) study. This may be explained by the similarity in final targets in the inflammatory pathway at which macrolides and NSAIDS appear to act. Meloxicam by inhibiting COX reduces the product of PGS (Burke et al., 2006), thereby causing vasodilatation (Vinay et al., 2005). On the other hand, macrolides inhibit the endothelial injury caused by oxygen-derived free radicals, and thus prevent an increase in vascular permeability, thus both the drug groups are affecting hemodynamic component of inflammation, albeit through different routes, and additional anti-inflammatory effect may therefore have not been seen by their combination.

## Author contributions

The original concept and study design was conceived by S.A. All experimental work and data analysis was performed by M.H., E.A. and M.Z. Figures were prepared by M.H., E.A. and M.Z. and the paper was written by M.H., E.A., M.Z, I.E., S.S and S.A.

## Acknowledgment

We would like to thank Dr. Ahmed Salahuldin and Dr. Amany I. El-Brairy for the gift of materials and some equipment required for experimentation. This work was partially funded by modern sciences and arts university, Giza, Egypt.

## Abbreviations

5-HETE: 5-Hydroxyeicosatetraenoic Acid
5-LOX: 5-Lipoxygenase Enzyme
FK506: Tacrolimus
FKBP: FK Binding Protein
IgG: Immunoglobulin G
IL: Interleukins
NSAIDs: Non-Steroidal Anti-Inflammatory Drugs
RA: Rheumatoid Arthritis
rRNA: ribosomal Ribonucleic Acid
TNF: Tumour Necrosis Factor
WBC: White Blood Cell

## References

Agen, A., Danesi, R., Blandizzi, C., Costa, M., Stacchini, B., Favini, P. (1993). Macrolide antibiotics as anti-inflammatory agents: Roxithromycin in an unexpected role. Agents and Actions 1-2:85–90.

Ahmed, N., Dawson, M., Smith, C. and Wood, E. (2007). Biology of Disease. 1st Ed, Taylor & Francis, New York, ISBN: 0748772103.

Banchroft, J.D., Stevens, A. and Turner, D.R. (1996). theory and practice of histoloicl techniques, 4th Ed, Churchil Livingstone, New york, London, San Francisco, Tokyo.

Barner A. (1996). Review of clinical trials and benefit/risk ratio of meloxicam. Scand. J. Rheumatol. Suppl. 102:29–37.

Barrett, K., Brooks, H., Boitano, S. and Barman, S. (2009). Ganong’s Review of Medical Physiology. 23rd Ed, McGraw-Hill Medical, New York, ISBN: 0071605673.

Burke, A., Smyth, E., FitzGerald, G.A. (2006). Analgesic-Antipyretic agents: Pharmacotherapy of Gout. 11th Ed, Goodman and Gilman’s The Pharmacological Basis of Therapeutics, New York.

Campagne, O., Mager, D. E., & Tornatore, K. M. (2019). Population Pharmacokinetics of Tacrolimus in Transplant Recipients: What Did We Learn About Sources of Interindividual Variabilities?. Journal of clinical pharmacology, 59(3), 309–325.

Chen, L., Deng, H., Cui, H., Fang, J., Zuo, Z., Deng, J., Li, Y., Wang, X., & Zhao, L. (2017). Inflammatory responses and inflammation-associated diseases in organs. Oncotarget, 9(6), 7204–7218.

Čulić, O., Eraković, V. and Parnham, M.J. (2001). Anti-inflammatory effects of macrolide antibiotics. European Journal of Pharmacology 429:209–229.

Cuzzocrea, S. (2005). Shock, inflammation and PARP. Pharmacological Research 52:72–82.

Davies, N.M. and Skjodt, N.M. (1999). Clinical pharmacokinetics of meloxicam: A cyclo-oxygenase-2 preferential nonsteroidal anti-inflammatory drug. Clin. Pharmacokinet 36:115–26.

Golan, D.E. (2008). Principles of pharmacology: the pathophysiologic basis of drug therapy. 2nd Ed, Lippincott Williams & Wilkins, Baltimore, ISBN: 0781783550.

Henderson, G., Dale, M.D., Ritter, J.M., Flower, R.J. and Rang, H.P. (2011). Rang and Dale’s pharmacology. 7th Ed, Elsevier/Churchill Livingstone, New York, ISBN: 0702034711.

Huang, M.T., Ghai, G. and Ho, C.T. (2004). Inflammatory process and molecular targets for anti-inflammatory nutraceuticals. Comprehensive reviews in food science and food safety 3:127–139.

Ianaro, A., Ialenti, A., Maffia, P., Sautebin, L., Rombola, L., Carnuccio, R., Iuvone, T., D’Acquisto, F., Di Rosa, M. (2000). Anti-inflammatory activity of macrolide antibiotics. J Pharmacol Exp Ther 1:156–163.

Johnson, M.D. (2011). Human Biology: Concepts and other issues. 6th Ed, Pearson Education, New York, ISBN: 0321701674.

Kannan, K., Kanabar, P., Schryer, D., Florin, T., Oh, E., Bahroos, N., Tenson, T., Weissman, J. S., & Mankin, A. S. (2014). The general mode of translation inhibition by macrolide antibiotics. Proceedings of the National Academy of Sciences of the United States of America, 111(45), 15958–15963.

Kelly, C., Chalmers, J. D., Crossingham, I., Relph, N., Felix, L. M., Evans, D. J., Milan, S. J., & Spencer, S. (2018). Macrolide antibiotics for bronchiectasis. The Cochrane database of systematic reviews, 3(3), CD012406.

Khobragade, A.A., Patel, S.B., Pophale, R.R. (2011). Analgesic and Anti-inflammatory Activity of Roxithromycin aand Erythromycin, Alone and in Combination with Ibuprofen: An Animal Study. IOSR Journal of Pharmacy 1:16–22.

Konno, S., Adachi, M., Asano, K., Okomoto, K. and Takahashi, T. (1993). Anti-allergic activity of roxithromycin: inhibition of interleukin-5 production from mouse T lymphocytes. Life Sci 52:25–30.

Lee, Y., Choi, J. Y., Fu, H., Harvey, C., Ravindran, S., Roush, W. R., Boothroyd, J. C., & Khosla, C. (2011). Chemistry and biology of macrolide antiparasitic agents. Journal of medicinal chemistry, 54(8), 2792–2804.

Majithia, V. and Geraci, S.A. (2007). Rheumatoid arthritis: diagnosis and management. Am. J. Med. 120:936–9.

Martina, B., Zeljko, K., Vesna, E., Michael, J. (2005). Cellular uptake and efflux of azithromycin, erythromycin, clarithromycin, telithromycin and cethromycin. Antimicrobial agents and Chemotherapy 1:2372–2377.

Maselli, D.J., Adams, S., Peters, J. (2011). The role of anti-invectives in the treatment of refractory asthma. Ther Adv Repir Dis. [Epub ahead of print].

Noble S. and Balfour J.A. (1996). Meloxicam. Drugs 5:424–30.

Patil, K. R., Mahajan, U. B., Unger, B. S., Goyal, S. N., Belemkar, S., Surana, S. J., Ojha, S., & Patil, C. R. (2019). Animal Models of Inflammation for Screening of Anti-inflammatory Drugs: Implications for the Discovery and Development of Phytopharmaceuticals. International journal of molecular sciences, 20(18), 4367.

Periti, P., Mazzei, T., Mini, E. and Novelli, A. (1992). Pharmacokinetic drug interactions of macrolides. Clin Pharmacokinet 23:106–131.

Porter, J. D., Watson, J., Roberts, L. R., Gill, S. K., Groves, H., Dhariwal, J., Almond, M. H., Wong, E., Walton, R. P., Jones, L. H., Tregoning, J., Kilty, I., Johnston, S. L., & Edwards, M. R. (2016). Identification of novel macrolides with antibacterial, anti-inflammatory and type I and III IFN-augmenting activity in airway epithelium. The Journal of antimicrobial chemotherapy, 71(10), 2767–2781.

Ransohoff, R.M. and Benveniste, E.N. (2006). Cytokines and the CNS. 2nd Ed, Taylor and Francis, Boca Raton, ISBN: 0849316227.

Rao, P.P.N., Kabir, S.N. and Mohamed, T. (2010). Nonsteroidal Anti Inflammatory Drugs (NSAIDs): Progress in Small Molecule Drug Development. Pharmaceuticals 3, 1530–1549.

Romero, L., Huerfano, C., & Grillo-Ardila, C. F. (2017). Macrolides for treatment of Haemophilus ducreyi infection in sexually active adults. The Cochrane database of systematic reviews, 12(12), CD012492.

Sasakawa, T., Sasakawa, Y., Ohkubo, Y. and Mutoh, S. (2004). FK506 ameliorates spontaneous locomotor activity in collagen-induced arthritis: implication of distinct effect from suppression of inflammation. International Immunopharmacology 5:503–510.

Serhan, C.N., Ward, P.A. and Gilroy, D.W. (2010). Fundamentals of Inflammation. 1st Ed, Cambridge university press, New York, ISBN: 0521887291.

Sevilla-Sánchez, D., Soy-Muner, D. and Soler-Porcar, N. (2010). Usefulness of Macrolides as Anti-inflammatories in Respiratory Diseases. Arch Bronconeumol. 46:244–254.

Shimizu, T., Kato, M., Mochizuki, H., Tokuyama, K., Morikawa, A. and Kuroume, T. (1994). Roxithromycin reduces the degree of bronchial hyperresponsiveness in children with asthma. Chest 106:458–461.

Tamaoki, J., Sakai, N., Tagaya, E. and Konno, K. (2000). Macrolide antibiotics protect against endotoxin-induced vascular leakage and neutrophil accumulation in rat trachea. Antimicrob Agents Chemother 38:1641–1643.

Trowbridge, H.O. and Emling, R.C. (1997). Inflammation: A Review of the Process. 5th Ed, Quintessence Publishing, Illinois, ISBN: 0867153105.

Vaz, A.P, Moris, A., Melo, N., Caetano Mota, P., Souto Moura, C., Amorim, A. (2011). Azithromycin as an adjuvant in cryptogenic organizing pneumonia. Rev Port Pneumol. 17:186–189.

Vázquez-Laslop, N., & Mankin, A. S. (2018). How Macrolide Antibiotics Work. Trends in biochemical sciences, 43(9), 668–684.

Vinay, K., Abul, K.A., Nelson, F. (2005). Acute and Chronic inflammation. 7th Ed, Pathological Basis of Disease, New Delhi: Elsevier.

Wallwork, B., Coman, M., Mackay-Sim, A., Greiff, L., Cervin, A. (2006). A double-blind, randomized, placebo controlled trial of macrolide in the treatment of chronic rhinosinusitis. Laryngoscope 2: 189–193.

White, M. (1999). Mediators of inflammation and the inflammatory process. J Allergy Clin Immunol 103:378–381.

Widmaier, E.P., Raff, H. and Strang, K. (2003). Vander’s Human Physiology: The Mechanisms of Body Function. 6th Ed, McGraw-Hill Higher Education, New York, ISBN: 0072880740.

Yao, C., & Narumiya, S. (2019). Prostaglandin-cytokine crosstalk in chronic inflammation. British journal of pharmacology, 176(3), 337–354.

Zappavigna, S., Cossu, A. M., Grimaldi, A., Bocchetti, M., Ferraro, G. A., Nicoletti, G. F., Filosa, R., & Caraglia, M. (2020). Anti-Inflammatory Drugs as Anticancer Agents. International journal of molecular sciences, 21(7), 2605.

